# Semi-supervised Omics Factor Analysis (SOFA) disentangles known and latent sources of variation in multi-omic data

**DOI:** 10.1101/2024.10.10.617527

**Authors:** Tümay Capraz, Harald Vöhringer, Klaus Sebastian Augusto Kruger Serrano, Ricardo Omar Ramirez Flores, Julio Saez-Rodriguez, Wolfgang Huber

## Abstract

A fundamental design pattern in biomolecular studies is to assay the same set of samples (organisms, tissue biopsies, or individual cells) by multiple different ‘omics assays. Group Factor Analysis (GFA) and its adaptation to high-dimensional settings, Multi-Omics Factor Analysis (MOFA), are widely used as a first-line approach to analyse such data and are effective in detecting patterns of correlation, organize them into so-called latent factors, and identify common and assay-specific factors. However, in many applications a subset of the found factors just rediscovers already known covariates (e.g., disease subtypes, environmental covariates) while others may represent genuine novelty.

Here, we present Semi-supervised Omics Factor Analysis (SOFA), a method that incorporates known covariates into the model upfront and focuses the factor discovery on novel sources of variation. We show SOFA’s effectiveness for discovering novel patterns by applying it to cancer, brain development and heart failure multi-omic data sets.

## Introduction

Using multiple “omic” measurement technologies on the same large set of biological specimens or samples is a fundamental design principle for studying biological variation^1–4^. With thousands or millions of features measured, the manifold hypothesis posits that the data concentrate along an (unknown) low-dimensional manifold inside the high-dimensional data space. Biological variation then corresponds to different positions on that manifold. Empirically, it has been found that useful coordinates on that manifold—what we might call interesting axes of biological variation—can be constructed by linear summary statistics. For instance, in principal component analysis (PCA) or its extension to factor analysis, each principal axis, or factor, is a linear combination of features, and this (weighted) averaging has the effect of reducing noise compared to looking at variation only by measuring individual features. Since typically a relatively small number of top principal axes or factors is considered, such approaches may also be viewed as dimension reduction techniques.

To take into account the multimodal structure of data, there are extensions of these methods including canonical correlation analysis (CCA)^5^, group factor analysis (GFA)^6^, multi-omics factor analysis (MOFA)^7^, iCLuster^8^, and MuVI^9^.

Applied to data, these methods return factors that, typically, fall into two categories: unsurprising and surprising. Unsurprising factors include those that align to known covariates (e.g., in the case of tissue biopsies, disease diagnosis, age or sex of the patient; in the case of environmental samples, geographical sampling location), or to technical artefacts^10^ (batch effects) associated with known experimental metadata. Surprising factors have no such easily identifiable explanation and are candidates for scientific novelty. The above-mentioned methods are unsupervised, in the sense that they do not use such already known information, and users are required to make their surprisingness considerations post-hoc^11^, in a process that is informal and error-prone.

Here, we present Semi-supervised Omics Factor Analysis (SOFA), a probabilistic factor model that jointly models multi-modal omics data and sample-level information (hereafter: guiding variables). SOFA approximates the data with a low-rank matrix decomposition and partitions the latent factors into guided factors each associated with a guiding variable, and unguided factors. In this way, already expected biological and technical drivers of variation can be regressed out from the latent space, making the interpretation of the remaining “surprising” factors simpler and potentially more fruitful.

Our model takes inspiration from prior work in the unimodal setting. Principal component regression (PCR) uses PCA factors as covariates in a regression model to predict guiding variables^12^. However, PCR does not guarantee that its covariates are relevant, since for their construction it only considers their covariance structure, but not the guiding variables. Partial least squares (PLS) simultaneously decomposes a high-dimensional set of predictors and guiding variables, aiming to find factors that capture the maximum covariance between the two^13,14^. SPEAR^15^ and DIABLO^16^ enable the incorporation of guiding variables of interest in the multimodal setting. Like PLS, these methods primarily uncover “unsurprising” factors that align with the guiding variables, rather than explicitly identifying “surprising” or unexpected factors.

We present four applications of SOFA that highlight its broad utility: to the pan-gynecologic cohort of The Cancer Genome Atlas (TCGA)^17^, to the cancer dependency map^18,19^ (DepMap), to single-cell multi-omic data of microglial cells^20^, and to a collection of data sets of acute and chronic human heart failure ^21–26^. We provide the SOFA method as an open source Python package along with comprehensive tutorials.

## Results

### The SOFA Model

The input to SOFA is a set of numeric data matrices containing multi-omic measurements from the same or mostly overlapping samples, and one or more covariates for each sample. The covariates can be continuous, binary or categorical and contain information that is known or suspected to be important for driving biological or technical variation between the samples. We also call these covariates *sample-level guiding variables*, and the different types of omics data *views*. From this input, SOFA extracts a lower-dimensional representation comprising a shared factor matrix and modality-specific loading matrices (**Fig. 1a**). SOFA partitions the latent factors into guided factors each associated with a guiding variable, and unguided factors (**Fig. 1b**). The loading weights of each factor represent, for each view, the molecular features that it depends on, and can usually be interpreted in terms of biological processes, gene expression programmes, modes of regulation, and the like (**Methods, Fig. 1c**). The factor matrix positions each sample in the latent space spanned by the factors, and each sample’s coordinates along the various factors can be interpreted as a degree of activity of a factor in the sample, similarly as in PCA or MOFA.

**Figure 1:**
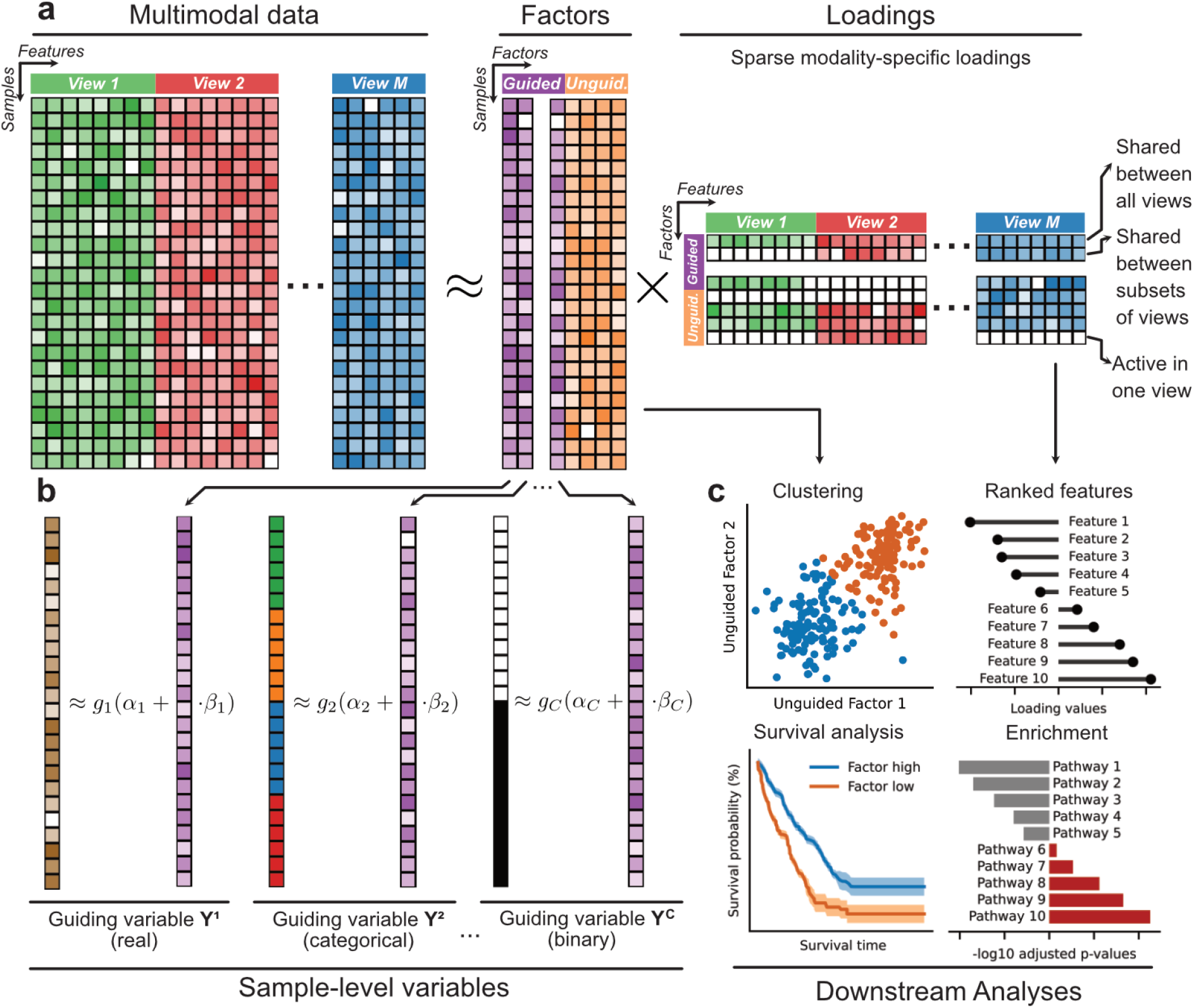
**a**: The SOFA model. Input data modalities X are decomposed into shared latent factors Z and modality specific feature loadings W. A hierarchical horseshoe prior shrinks the loading weights on the view, factor and feature level. This leads to factors shared between all views, factors active in one view and factors shared between subsets of views. **b:** Guided factors are linked to the guide variables. **c:** Factors not linked to a guide variable capture potentially novel sources of variation. Factors and loadings can be used for various sample- and feature-level downstream analyses.

SOFA is a hierarchical Bayesian factor model, and we ensured scalability of its inference to large multi-omic data sets by employing stochastic variational inference (**Methods**). Our inference method for SOFA employs a sparsity inducing prior^27^, which pulls the loading weights of some factors for some modalities to zero, if the data are not pulling them elsewhere strongly enough. As a result, we can distinguish between variation that is commonly shared across modalities and variation that is present in only one or a few modalities.

### SOFA identifies cancer type independent immune infiltration vs proliferation axis in the TCGA pan-gynecologic data set

We applied SOFA to a pan-gynecologic cancer multi-omic data set of the cancer genome atlas (TCGA) project, which profiled the transcriptome (mRNA), proteome, methylome, and miRNA of 2599 samples from five different cancers^17,28–31^. Additionally, the study includes data about mutations, metadata, and clinical endpoints progression free interval (PFI) and overall survival (OS).

We performed a SOFA rank-12 decomposition, using five guided factors with binary labels for each cancer type and seven unguided factors. Guided factors showed a clear separation based on the binary cancer type labels (**Fig. 2a**). We then verified that the unguided factors did not capture cancer type-related variance by computing Adjusted Rand Index (ARI) scores to assess clustering alignment with cancer labels (**Fig S2e, Methods**). This analysis showed that the set of unguided factors had the lowest alignment with cancer type labels, suggesting that the guided factors mostly explained the cancer-associated variability. Of note, ARI scores for the unguided SOFA decompositions at both rank-7 and rank-12 exhibited lower cluster alignment in comparison to the guided factorisation, indicating that factor guidance helped disentangle cancer-associated variability (**Fig. S2a**,**c**,**e**).

**Figure 2:**
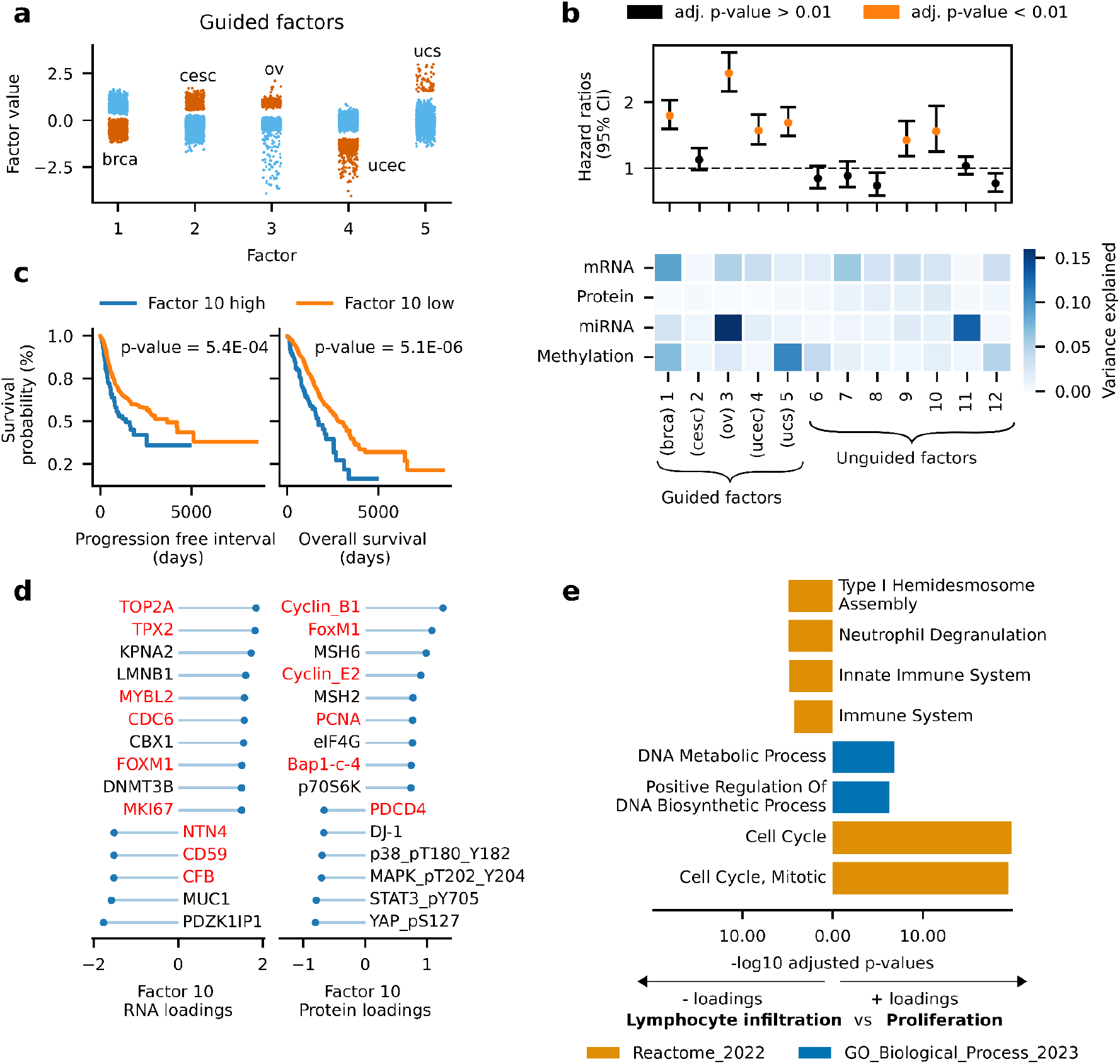
**a:** Distributions of guided factors colored by their guiding variables (orange). **b**: Top: Hazard ratios from univariate Cox regression for each factor on overall survival. Hazard ratios > 1 indicate that positive factor values are associated with higher risk. Hazard ratios are colored if adjusted p-values are lower than 0.01. P-values were adjusted for multiple testing using the Benjamini-Hochberg (BH) method. Bottom: Variance explained by each factor for each modality. **c**: Kaplan-Meier curves showing progression free survival (left) and overall survival (right) for samples with factor values < 50th and > 95th percentiles of Factor 10. **d**: Features with the highest absolute loading weights for factor 10 in transcriptomics (left) and proteomics loadings (right). **e**: Gene set overrepresentation analysis for the transcriptomics features with the highest absolute loadings of factor 10. The −log10 adjusted p-values are multiplied by the sign of the loadings to reflect the direction of the association with the factor. Cancer types shown: brca: breast invasive carcinoma; cesc: Cervical squamous cell carcinoma and endocervical adenocarcinoma; ov: Ovarian serous cystadenocarcinoma; ucec: Uterine Corpus Endometrial Carcinoma; ucs: Uterine Carcinosarcoma

To evaluate the importance of each factor, we calculated the fraction of variance explained by each factor across the input modalities (**Fig. 2b**). For example, Factor 3 explained the highest fraction of variance in the miRNA data, consistent with previous research suggesting miRNA expression is critical in ovarian cancers and may serve as a biomarker^32,33^. Factor 1, representative for breast cancers, showed breast cancer associated proteins AR^34^, PR^35^ and ER-alpha^36^ among the largest proteomics loading weights (**Fig. S2f**). Factor 3 (ovarian serous cystadenocarcinoma) featured *BCAM*^*37*^, *MALAT1*^*38*^ *and WT1*^*39*^ among the highest transcriptomics loadings weights (**Fig. S2g**).

We then explored the SOFA factors for potential biological relevance by testing them for association with disease outcome, using univariate Cox regression of each factor on overall survival (OS, **Fig 2b**). Notably, Factor 3, linked to ovarian serous cystadenocarcinoma, was significantly linked to poor survival (Hazard Ratio (HR)=2.44 95% CI: 2.17-2.75). Two out of seven unguided factors showed significant associations with OS, suggesting that most of these factors do not strongly influence survival outcomes. Among the unguided factors predictive of survival, we further investigated Factor 10, which had the highest HR among the unguided Factors (HR=1.56 95% CI: 1.25-1.94, **Fig. 2b**). Stratifying samples by Factor 10 values, followed by non-parametric survival curve estimation, revealed substantial differences in median survival times between groups (OS: Median survival time 1701 vs. 2888 days, PFI: 1396 vs 3669 days, **Fig. 2c**). Negative loadings of Factor 10 featured *NTN4, CD59* and *CFB*, and proteins including PDCD4, indicative of complement system activation and lymphocyte infiltration^40–43^(**Fig. 2d**). In contrast, positive loadings included proliferation markers *TOP2A* and *TPX2*, as well as Cyclin B1, FoxM1^44–47^, suggesting a signature of increased proliferation. This was further corroborated by performing a gene set overrepresentation analysis on the top transcriptomics loadings. Negative loadings were associated with immune system related gene sets, while those with positive loadings were enriched in cell cycle and proliferation related terms (**Fig. 2e**). Finally, we note that Factor 10 was not restricted to a single cancer type (**Fig. S2b,d**), indicating that it captured biological signatures common to all cancers. Taken together, we found that the guided SOFA factorization delineated cancer type-specific variation from broader biological processes, such cell proliferation or immune cell infiltration, thereby facilitating their identification and subsequent downstream analyses.

### SOFA identifies EGFR dependency axis in pan-cancer dependency map, while accounting for known driver mutations and cell growth

Next, we analyzed a pan-cancer cell line data set of the Cancer Dependency Map (DepMap)^19,48^ together with proteomics data from the ProCan-DepMapSanger study^18^. The DepMap project aims to identify cancer vulnerabilities and drug targets across a diverse range of cancer types. The data set includes multi-omic data, encompassing transcriptomics^49^, proteomics^18^, and methylation^50^, as well as drug response profiles for 627 drugs^50^, and CRISPR-Cas9 gene essentiality scores^48^ for 17,485 genes for 949 cancer cell lines across 26 different tissues.

To identify CRISPR-Cas9 gene essentiality scores associated with multi-omic features, we performed a rank-20 SOFA decomposition to the DepMap data, while accounting for potential/known drivers of variation such as growth rate, microsatellite instability (MSI) status, *BRAF, TP53* and *PIK3CA* mutation and hematopoietic lineage. Although CRISPR-Cas9 essentiality scores could technically be included as an additional view in the SOFA decomposition, we chose to exclude them to enable independent testing of factor associations with these scores.

A t-SNE of all the inferred factors showed that they captured the differences between cancer types (**Fig. 3a**). Further, we confirmed successful factor guidance by asserting that SOFA extracted high or low factor values in a variable-dependent manner (**Fig. 3c**). We then focused on the guided factors and observed that Factor 6, consistent with previous studies identifying hematopoietic lineage as a major driver of variation in cell lines^18^, explained the largest fraction of variance (34%) across all data modalities (**Fig. 3b**). Factor 1, guided by growth rate, mostly explained variance in drug response (**Fig. 3b**), which is expected as the natural logarithm of the half maximal inhibitory concentration (lnIC50) drug response is derived from the growth rate of the cell line. We then subjected the loadings of guided factors to gene set overrepresentation analysis. Expectedly, Factor 3, guided with labels for BRAF mutation, a common mutation in skin cancer^51^, was associated with skin cancer related gene sets (**Fig. S3c**). The loadings of Factor 6, guided with hematopoietic lineage, displayed gene sets related to the immune system (**Fig. S3d**). Statistical testing revealed that Factors 3 and 4 were significantly associated with CRISPR-Cas9 essentiality scores of BRAF and TP53, the mutations used to guide these factors (**Fig. S3e-f, Methods**).

**Figure 3:**
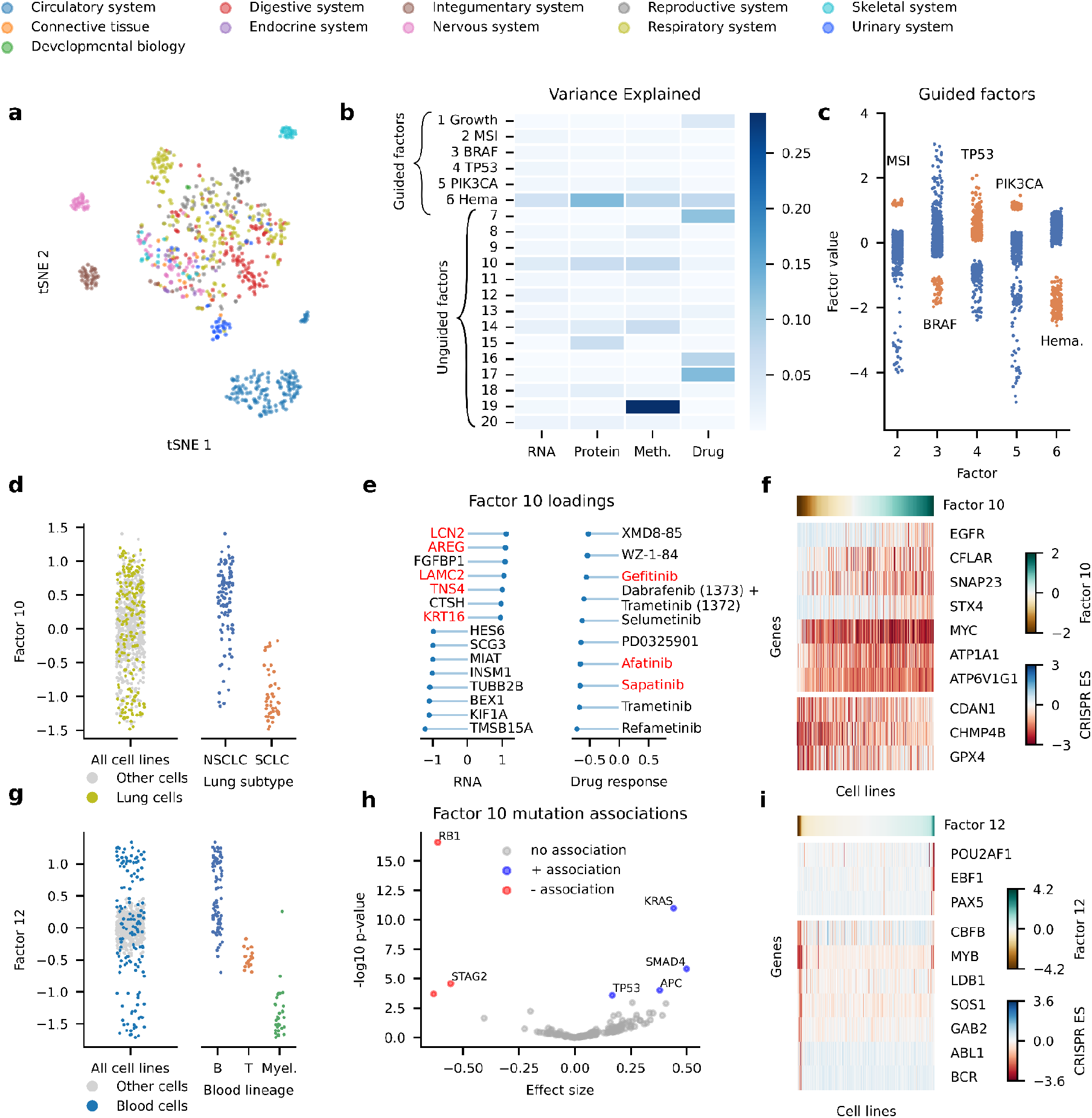
**a:** tSNE of all SOFA factors, colored by cancer types. **b**: Variance explained by each factor for each modality. **c**: Distributions of guided factors colored by their guiding variables (orange). **d**: Distribution of Factor 10 cell lines from the respiratory system are colored (left). Distribution of Factor 10 for cell lines from non-small cell lung cancer (NSCLC) and small cell lung cancer (SCLC) (right). **e**: Highest absolute transcriptomics (left) and drug response (right) loadings of Factor 10. **f**: Heatmap showing CRISPR essentiality scores (ES) of top 10 genes significantly associated with Factor 10, tested using a linear model. The cell lines are ordered based on their values of Factor 10. **g**: Distribution of Factor 12, cell lines from the circulatory system are colored (left). Distribution of Factor 12 for B-cell, T-cell and myeloid cell lines (right). **h**: Volcano plot of associations between Factor 10 and mutations, tested using a two sample t-test. BH-adjusted p-values < 0.05 are highlighted in color. Genes colored in blue are significantly associated with higher and genes colored in red with lower factor values. **i**: Heatmap showing CRISPR essentiality scores (ES) of top 10 genes significantly associated with Factor 12, tested using a linear model. The cell lines are ordered based on their values of Factor 12.

Among the unguided factors, Factor 10 had the highest values in colorectal and lung cancer cell lines (**Fig. S3a**). A closer examination revealed that this factor delineated non-small cell lung cancer (NSCLC) from small cell lung cancer (SCLC) (**Fig. 3d**). Top-ranking genes featured regulators of the epidermal growth factor receptor (EGFR), including *LCN2, AREG, LAMC2, TNS4* and *KRT16*^*52–56*^ (**Fig. 3e**). Similarly, the lowest loadings for drug response lnIC50 values, indicating sensitivity, were EGFR targeting drugs Gefitinib, Afatinib and Sapatinib (**Fig. 3e**). To confirm the association between the EGFR pathway and Factor 10, we tested for associations with CRISPR-Cas9 essentiality scores, indeed EGFR was a top ranking association (**Fig. 3f**). Further, high Factor 10 values were associated with cell lines harboring *EGFR, TP53, APC, SMAD* and *KRAS* mutations, which were previously described to play a role in progression of colorectal cancer^57,58^ (**Fig. 3h**).

Factor 12 separated B cells, T cells, and myeloid cells within the hematopoietic lineage (**Fig. 3g** and **S3b**). Correlating Factor 12 with CRISPR-Cas9 essentiality scores showed that positive factor values (B-cells) were significantly associated with B-cell markers *EBF1, PAX5* and *MEF2B*^*59,60*^, while negative values (myeloid cells) were significantly linked to myeloid markers *MYB* and *CBFB*^*61,62*^ (**Fig. 3i**). In summary, by leveraging multi-omic data together with CRISPR-Cas9 essentiality scores, we identified an unguided factor linked to the EGFR pathway, along with another factor that explained the difference between hematopoietic cell types.

### SOFA disentangles between and within cell type variation in a single-cell multi-omic data set of the human developing cerebral cortex

We applied SOFA to infer 15 factors from data of a single-cell multi-omic experiment^20^, where the authors simultaneously profiled the transcriptome (RNA) and chromatin accessibility (ATAC) of 45,549 single cells of the human cerebral cortex at six different developmental stages. The authors identified 13 different cell types in the data.

We assumed that guiding a subset of SOFA factors with cell type labels could help reveal variations within cell types through the unguided factors. To test this, we conducted a rank-15 SOFA decomposition on the single-cell multi-omic data set, guiding the first 13 factors with binary cell type labels while leaving the remaining two factors unguided to capture sources of variation within cell types.

**Fig. 4a** shows the fraction of variance that each factor explained in the RNA and ATAC modalities, with factors guided by microglial and oligodendrocyte labels explaining most variance. The UMAP embedding of all factors in **Fig. 4b** showed that the cells cluster by cell type clearly separating the different cell types.

**Figure 4:**
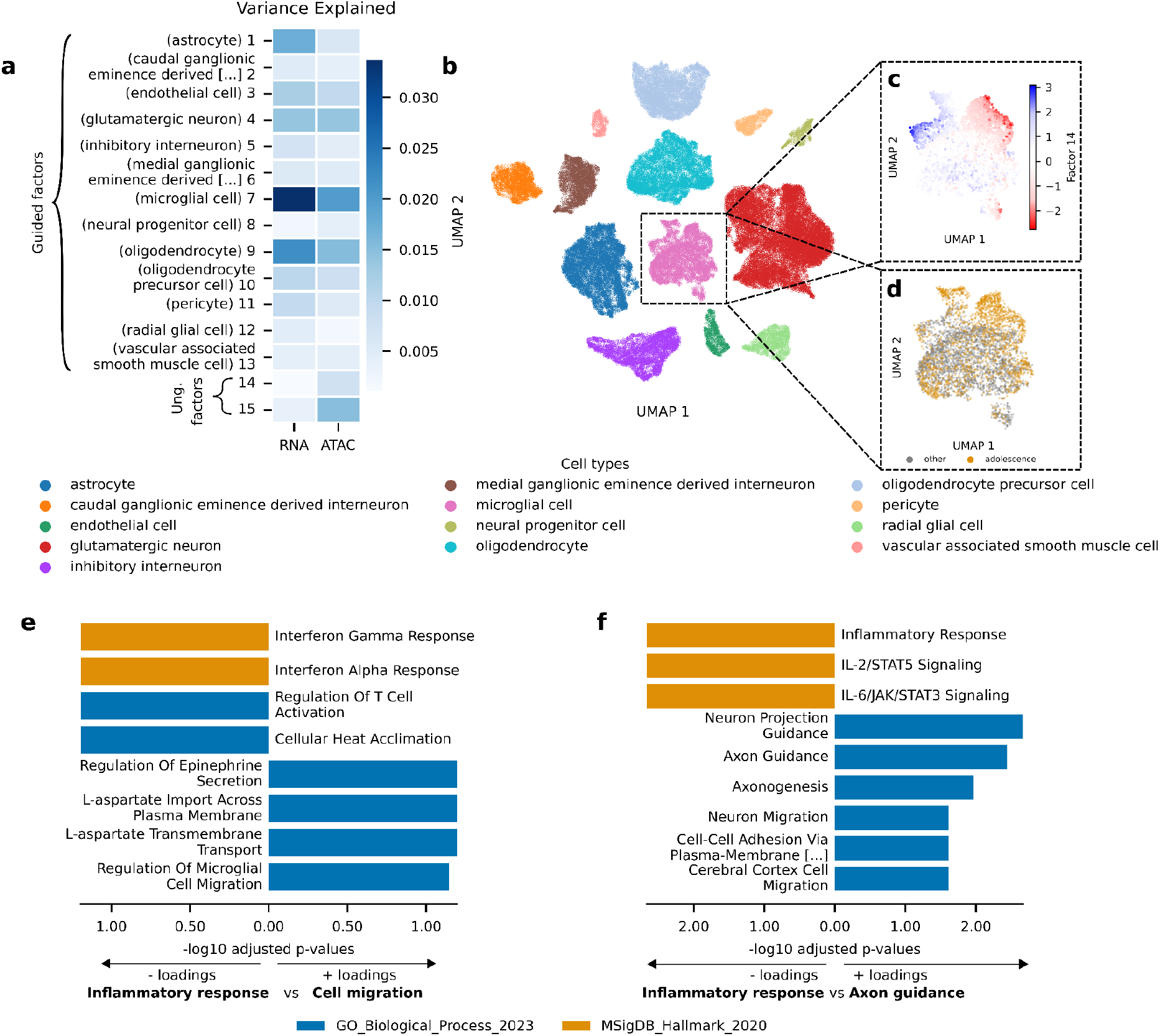
**a**: Variance explained in RNA and ATAC data modality by each factor. **b**: UMAP based on all factors coloured by cell type. **c**: UMAP of microglial cells coloured by Factor 14. **d**: UMAP of microglial cells colored by adolescence age group. **e**: Gene set overrepresentation analysis for the transcriptomics features with the highest absolute loadings of factor 14. The −log10 adjusted p-values are multiplied by the sign of the loadings to reflect the direction of the association with the factor. **f**: Gene set overrepresentation analysis for the ATAC features with the highest absolute loadings of factor 14. The −log10 adjusted p-values are multiplied by the sign of the loadings to reflect the direction of the association with the factor.

First, we checked whether the loadings of the guided factors picked up biological signals related to their guiding covariate. Indeed, the guided factors separated cells based on their guiding cell type labels (**Fig. S4a**). The top loadings of the guided factors were cell type specific markers of their respective guiding cell type label. Highest transcriptomics loadings of Factor 1 (astrocytes) included astrocyte specific marker genes such as *SLC1A2*, SLC1A3^63^ and *GLUL*^*64*^. Factor 7 (microglial cells) captured microglia markers *CSF1R*^*65*^, *TYROBP*^*66*^ and *CX3CR1*^*67*^. Factor 9 (oligodendrocytes) had *MBP*^*68*^, *MOBP*^*69*^ and *PLP1*^*70*^ among the top transcriptomics loadings.(**Fig. S4b-d**).

When further inspecting the unguided factors, we found that Factor 14 captured variation within microglial cells and was only active in a subset of microglial cells with relatively high and low factor values (**Fig. 4c**). Gene set overrepresentation analysis on the transcriptomics and ATAC loadings of Factor 14, revealed that the positive loadings are related to microglial migration and transport processes, while the negative loadings play a role in inflammatory response and interleukin signaling (**Fig. 4e** and **4f**). Previous studies using single-cell RNA-seq data from microglia indicated that there is a diverse range of microglial subtypes^71^, including populations expressing genes related to inflammatory response^72^ and migration^71^. Furthermore, microglia play a critical role in brain development^73^, especially during adolescence^74^ and Factor 14 was mostly active in cells from the adolescence age group (**Fig. 4c-d**).

This shows that SOFA is also capable of finding biologically interesting patterns, such as within cell type variation, in single-cell multi-omic data sets.

### Comparison of SOFA and MOFA in the analysis of heart failure single-cell data

We compared SOFA’s guided factor analysis approach to MOFA’s unsupervised one to evaluate whether SOFA enhances detection of covariate-associated low variance signals, while preserving the integrity of factors representing dominant variability. We applied both methods in the context of multicellular integration of single-cell RNA-Seq data, that aims at inferring coordinated gene expression programs between cell-types in tissues, across a previously curated collection of five datasets of acute and chronic human heart failure ^21–26^. This collection of data sets includes single-cell RNA-Seq data of seven cell types from chronic human heart failure cases (HF) and controls (NF), along with covariates including age, sex, and heart failure status, a binary variable indicating patients with any heart disease etiology that required heart transplantation (**Fig. S5a-b**). To apply SOFA and MOFA, we pseudo bulked the gene expression of each cell type and represented them as separate views (**Fig. S5c**). This enables the inferred factors to capture gene expression coordination across cell types—referred to as multicellular programs—which could then be associated with patient-level covariates. Heart failure was expected to be the primary source of variability in these data sets^21^. However, given the smaller cohort sizes compared to our previous analysis examples (maximum size = 79), low variance covariates like age and sex might be challenging to detect using an unsupervised model. We fitted a MOFA model and two SOFA models: one guided by age and sex, and another that included guidance from both age and sex along with heart failure binary status. This approach allowed us to assess whether SOFA could identify factors associated with less dominant covariates and evaluate the impact of factor guidance on the detection of unguided factors representing major sources of variability, in this case heart failure. We observed that both methods recovered a comparable fraction of variance associated with heart failure and age. However, SOFA models recovered a greater fraction of variance (mean=3.65% SOFA HF guided; mean=3.7% SOFA HF unguided) associated with sex across data sets compared to MOFA (mean=0.43%), which showed near-zero values in all but one data set (**Fig. S5d**). To assess whether guiding SOFA with known covariates would hamper the unguided factors to capture the main sources of variability — here heart failure —, we correlated the heart failure-associated factors between the MOFA model and the SOFA model not guided with heart failure. The mean Spearman correlation of 0.81 and 0.870 at both the factor score and gene loading levels indicated minimal influence of factor guidance on these unguided factors. Next, we evaluated whether the loadings of the heart failure and sex-associated factors across the three models captured similar results as the ones expected from differential expression analyses. First, we used DEseq2 models to identify differentially expressed genes between failing and non-failing hearts, and males and females, across cell-types for each data set independently (adj. P-value < 0.05). Then we sampled an identical number of genes from the gene loadings ranked by their absolute value. We observed higher recall for heart failure of the SOFA model guided with heart failure (mean=66.2%) compared to the other two models (mean=55.05% SOFA HF unguided; mean=55.25% MOFA) (**Fig. S5e**). For the sex-associated factors we observed a mean recall of 55.84% in the SOFA models. Together, these results highlight that guided SOFA factors can enhance the detection of gene expression profiles associated with low variance covariates like sex while preserving the ability to infer major sources of variability, providing a flexible alternative to fully unsupervised models.

## Discussion

SOFA is a fully interpretable probabilistic framework for the analysis of multi-omic data.

It is based on group factor analysis methods such as GFA and MOFA^6,7^ and decomposes multi-omic data into latent factors that capture shared or modality-specific variance across multiple omics modalities. In contrast to existing latent factor models, it enables the incorporation of a priori known (or suspected) drivers of variation. The drivers of variation can be covariates such as mutations, pathways, cancer or cell types. While the guided factors explain variation in the molecular omics data that is driven by the guiding variables, the unguided factors can capture the remaining variation, potentially leading to novel biological insights.

Previously reported analysis approaches to such data either involved manual post-hoc annotation of factors, or splitting of the data into subsets of cell types, tissues etc., to identify subset specific factors^18,21,75^. SOFA offers a potentially easier and more straightforward workflow, by employing the complete data and the guiding variables in a single end-to-end analysis.

The versatility of SOFA was showcased through four examples of exploratory data analysis. Analysis of the pan-gynecologic cohort of the TCGA^17^ revealed a biological stratification predictive of survival and independent of cancer type. In a second example, we identified a latent factor related to EGFR cancer dependency enriched in lung and colorectal cancer samples in a pan-cancer cell line analysis of the DepMap^18,19^. Applying SOFA to a single-cell multi-omic data set from cells of the developing human cerebral cortex^20^, we show that SOFA can account for inter cell type heterogeneity, while discovering factors explaining intra cell type heterogeneity. We compared SOFA and MOFA, a method that does not offer the ability to guide factors with known covariates, in an integrative analysis of five data sets of acute and chronic human heart failure SOFA is a tool for exploratory analysis of multi-omic data sets. It identifies sets of features (e.g., representing genes and gene products) that covary across multiple modalities. It is aimed to generate hypotheses that will be validated with independent test data or follow-up experiments. SOFA is a linear method, which facilitates interpretation, but would be a limitation if there are important phenomena that are only evident through non-linear functions of the features. The current implementation of SOFA considers each omics view with a Gaussian likelihood. For highly heteroskedastic data, such as, for instance, RNA-seq, appropriate preprocessing and transformation is required.

In summary, SOFA is able to discover novel axes of variation while accounting for previously known variation. It is implemented using the probabilistic programming framework Pyro^76^, is available as a Python package including documentation and analysis examples, and its processing speed makes it amenable to interactive analysis on data sets such those shown here.

## Methods

### The SOFA model

SOFA jointly decomposes a multimodal data set 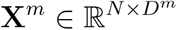 with *m =* 1,…,*M* modalities for *i =* 1 …*N* samples and *i =* 1,…,*D*^*m*^ features, and a set of continuous or categorical sample-level covariates (hereafter guiding variables) **Y**^1^,…, **Y**^*C*^ where **Y**^*C*^ ∈ ℝ^*N*^ or 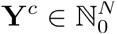 with guiding variables into a lower dimensional shared latent factor matrix and modality specific loading matrices. We model the entries 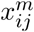 of **X**^*m*^ as normally distributed, given an unobserved mean expression level μ_*ij*_ and a feature-specific scale parameter 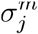, and the entries 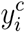 of with a mean/location parameter 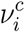 and an optional scale parameter *k*^*c*^

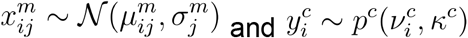

where *p*^*c*^ denotes an appropriate likelihood (i.e. Bernoulli, Categorical or Normal) for the ‘th guiding variable. In SOFA we express 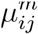 and 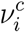

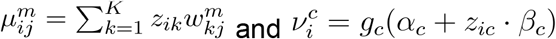

where 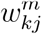 represents the entries of the *m*th loading weight matrix 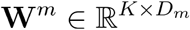 consisting of *k* = 1,…,*K* components where *C* ≤ *K* ≤min(*D*_1_,…, *D*_*M*_), *z*_*ik*_ the elements of a factor weight matrix **Z** ∈ ℝ^*N* × *K*^ containing the coordinates of each sample with respect to **W**^*m*^, *α*_*c*_ and *β*_*c*_ an offset and slope parameter and *g*_*c*_ an appropriate inverse link function for the ‘th guiding variable.

SOFA is formulated as a Bayesian probabilistic model and uses prior distributions on the model parameters (for a complete derivation of the model see the Supplementary Methods).

### Interpretation of the factors

Analogous to the interpretation of factors in PCA, SOFA factors ordinate samples along a zero-centered axis, where samples with opposing signs exhibit contrasting phenotypes along the inferred axis of variation, and the absolute value of the factor indicates the strength of the phenotype. Importantly, SOFA partitions the factors of the low-rank decomposition into guided and unguided factors: the guided factors are linked to specific guiding variables, while the unguided factors capture global, yet unexplained, sources of variability in the data.

### Interpretation of the loading weights

SOFA’s loading weights indicate the importance of each feature for its respective factor, thereby enabling the interpretation of SOFA factors. Loading weights close to zero indicate that a feature has little to no importance for the respective factor, while large magnitudes suggest strong relevance. The sign of the loading weight aligns with its corresponding factor, meaning that positive loading weights indicate higher feature levels in samples with positive factor values, and negative loading weights indicate higher feature levels in samples with negative factor values.

### Parameter inference

SOFA is formulated as a Bayesian probabilistic model and uses black box variational inference^77^ to infer the unobserved parameters. Black box variational inference uses a simpler variational distribution to approximate the intractable posterior. It is a scalable technique that uses stochastic optimization to optimize the evidence lower bound (ELBO) with respect to the variational parameters of the variational distribution. The ELBO is optimized using stochastic variational inference by taking Monte Carlo samples from the variational distribution and computing unbiased but noisy gradients with respect to the variational parameters from the ELBO^77,78^. The ELBO can be used to assess convergence.

For the implementation of SOFA we used the probabilistic programming framework Pyro^76^. If necessary, e.g. for single-cell data, we compute the gradients over subsamples of batches of the observations.

### Model selection

The number of factors is an important hyperparameter of SOFA that needs to be determined by running the inference multiple times with a varying number of factors. Factors typically become correlated if the decomposition rank is too high. Additionally, RMSE and variance explained can be used to assess good choices of the number of factors.

Furthermore, guiding variables potentially driving variation in the data, need to be tested in multiple inference runs.

### Simulations

To verify that SOFA is able to both infer factors that approximate the multi-omic data and simultaneously guide a subset of the factor, we simulated data from the generative model, inferred a varying number of latent factors and computed the root mean square error between the reconstructed and the simulated data.

Fig. **S1a** shows the RMSE for the simulated multi-omic data for an unguided and a SOFA model where the first two factors were guided with simulated guide variables. The RMSE of the unguided model was slightly lower than that of the guided model. However, this was only the case for model fits with a lower number of factors than the simulated data. This indicates that if the number of latent factors is at least equal to the number of latent dimensions of the simulated data, SOFA is able to guide a subset of the factors without losing its ability to fit the multi-omic data. Furthermore, the simulated guide variables align with their respective guided factors (**Fig. S1b,c**).

### Comparison of SOFA and MOFA in the analysis of heart failure single-cell data

To investigate how guiding a subset of factors influences factorization, we applied SOFA for multicellular integration^21^ to a collection of single-nucleus RNA-Seq data sets of acute and chronic human heart failure compiled in^22^. For each study, additionally, patient covariates such as age, sex and heart failure labels were available. Cell type annotations of each single-nucleus data were aligned to a unified ontology of seven cell-types: Cardiomyocytes, fibroblasts, pericytes, and endothelial, vascular smooth muscle, myeloid and lymphoid cells.

Pseudobulk expression profiles of each cell type per patient were used as input views for SOFA, as outlined in^21^. Briefly, pseudobulks for each cell type were included only if they contained at least 20 unique cells. Cell types present in fewer than 40% of the patients in a given data set were removed. At the gene level, genes expressed fewer than 20 times across 60% of the patients in a cell type were considered lowly expressed and filtered out. After this step, cell types with fewer than 50 expressed genes were discarded. Samples with less than 97% coverage of the filtered gene set were also excluded. Known background genes were removed, retaining only genes of interest for factor inference. Following these filtering steps, only cell types with at least 50 remaining genes were included in the final analysis.

We fit three models: a rank-10 MOFA model, a rank-10 SOFA model guided by age and sex, and a rank-10 SOFA model guided by age, sex, and heart failure. SOFA models were fitted with a total of 3500 steps and a learning rate of 0.01.

We identified factors associated with heart failure, sex, and age, and calculated the fraction of variance explained by these factors. This metric served as a proxy for the amount of variation in the omics data linked to heart failure, sex and age. Analysis of variance was used to associate factor scores with heart failure and sex, and Pearson correlation was used to test for association with age. When multiple factors associated with a covariate, we summed their fraction of variance explained.

To identify the most similar pair of heart failure-associated factors between the MOFA model and the SOFA model guided by age and sex in each study, we used absolute Spearman correlations of gene weights and factor scores, together with their fraction of explained variance. For each data set and model, we identified genes significantly associated with heart failure (adjusted p-value <0.05), using the R package DESeq^79^ with default parameters to filter out lowly expressed genes. We selected the number of significant genes among the highly ranked genes in the model by using their absolute factor loading as the basis for ranking. We then calculated recall, defined as the percentage of loadings that included these expected genes in the right regulatory direction.

### Data processing

All data processing steps were performed using the *muon* (version 0.1.3)^80^ and *anndata* (version 0.8.0)^81^ Python packages.

### TCGA pan-gynecologic atlas

All data modalities, survival data and metadata were downloaded using the R package *curatedTCGAdata* (version 1.26.0)^17,82^. All data modalities except the protein array data were log-transformed. All data was centered and scaled to unit variance. Before the analysis, we selected the top 2000 highly variable features of the RNA-seq and methylation data sets.

### Pan-cancer DepMap

We analyzed a multi-omic cancer cell line data set with proteomic data from Gonçalves et al.^18^ combined with multi-omic and CRISPR essentiality data^19,48^ of the same cell lines. Processed and transcript per million (TPM) normalized RNA-seq data^49^, mutation data^50^ and integrated CRISPR essentiality scores were obtained from https://cellmodelpassports.sanger.ac.uk/^83^. Processed methylation beta values^50^ were obtained from https://www.cancerrxgene.org. Processed and normalized proteomics abundances and drug response lnIC50 data^50^ was obtained via figshare from Gonçalves et al.^18^. We log-transformed, centered and scaled the RNA-seq TPM data and the pre-processed methylation beta values and selected the top 2000 highly variable genes prior to the analysis. The proteomics data was imputed with zero, and the top 2000 highly variable proteins were selected. Additionally, we regressed out the sample mean as suggested in Gonçalves et al.^18^. The drug response lnIC50 values were centered and scaled.

### Single-cell multi-omics of the human cerebral cortex

The single-cell RNA-seq and ATAC-seq data was obtained from Zhu et al.^20^ and downloaded in the h5ad format from https://cellxgene.cziscience.com/. RNA-seq data was normalized to library size and log-transformed. We selected the top 2000 highly variable genes in both modalities. Both data sets were centered and scaled before analysis. Cell type labels were obtained from the h5ad objects.

### Downstream analysis

#### Association testing

Associations between factors and mutations were tested using two sample t-tests.

To test for associations between the guided SOFA factors and essentiality scores, we used factor values as predictors in a linear regression model to explain essentiality scores, and then evaluated the significance of the fitted regression coefficient. Briefly, Genes with highly negative CRISPR-Cas9 essentiality scores are deemed essential, as their loss significantly impairs cell viability. Scores near zero indicate nonessential genes with minimal impact on viability, while positive scores suggest that gene loss may confer a survival advantage. Therefore, negative coefficients (effect sizes) indicate that higher factor values correspond to lower essentiality scores, implying a decrease in cell viability. All p-values were adjusted for multiple testing using the Benjamini-Hochberg procedure^84^.

#### Survival analysis

The survival analysis was performed using the *lifelines* package (version 0.27.7)^85^. Assessment of whether a factor was predictive of overall survival was done by per factor univariate Cox’s proportional hazard models. Resulting p-values were adjusted for multiple testing using the Benjamini-Hochberg procedure ^84^.

#### Gene set overrepresentation analysis (ORA)

To annotate the factors with biological processes we used the enrichr API^86^ provided in the *gseapy* Python package (version 1.0.4) ^87^. We used the top 100 features with highest or lowest loadings of a given factor and performed gene set overrepresentation analysis using all genes in the data set as background.

#### Fraction of explained variance

Assuming the data **X** is centered, the fraction of variance that is explained by factor *k* of modality *m* was calculated as follows:

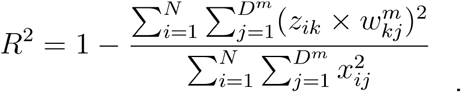

Where the denominator corresponds to the total sum of squares of the centered data **X** and the nominator to the residual sum of squares. The fraction can be understood as the variability a given factor *k* accounts for.

#### Two dimensional visualizations of the latent space

The t-distributed Stochastic Neighbor Embeddings (tSNE) for 2D visualizations of the latent space were calculated using the *scikit-learn* Python package (version 1.2.1)^88^. Uniform Manifold Approximation and Projection (UMAP)^89^ embeddings were computed using the implementation provided in the *scanpy* Python package (version 1.9.3) ^90^.

#### Root mean square error for model selection

To assess the quality of the individual model fits we calculated the root mean square error as follows:

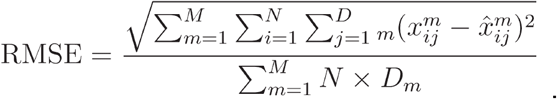

#### Adjusted Rand Index

To verify that the unguided factors did not capture variance associated with categorical guiding variables, such as cancer type labels, we compared the Adjusted Rand Index (ARI)^91^ for clusterings on the latent factors of a guided vs unguided model. To this end, we used the *scib* package^92^ to perform clustering on the latent factors and calculate the ARI between the cluster labels and categorical guiding variables for 10 independent initializations. The ARI measures how well the clusterings align with the guiding variables; a higher ARI corresponds to greater overlap, thus indicating that the latent factors captured variance associated with the guiding variables.

## Software availability

SOFA is implemented in the Python package *biosofa* and is available on github (https://github.com/tcapraz/SOFA) and PyPI (https://pypi.org/project/biosofa/), along with documentation and example analysis notebooks. Code and processed data to reproduce the analysis and figures are available on Zenodo https://doi.org/10.5281/zenodo.14761127.

## Acknowledgements

We would like to thank Stefan Peidli and Martin Rohbeck for valuable feedback and discussions. We thank EMBL IT services and HPC resources for providing infrastructure to perform computations needed for this work. The results shown here are in part based upon data generated by the TCGA Research Network: https://www.cancer.gov/tcga.

## Supplementary Figures

**Figure S1:**
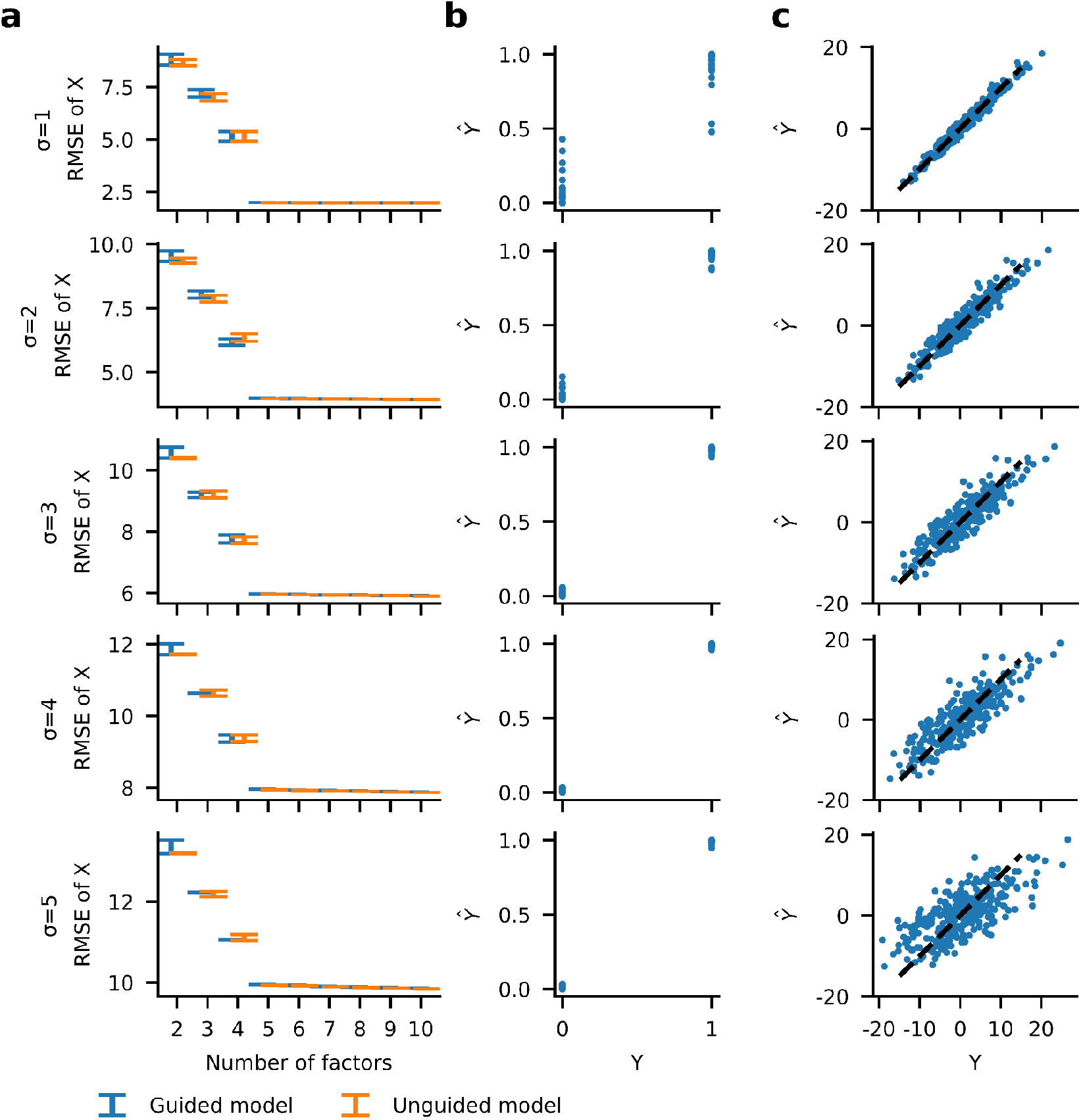
In-silico data were simulated according to the generative model of SOFA using a true rank of 5 and two guiding variables for the first (binary) and second factor (continuous). The simulated multi omic data comprised 2 views with each 300 observations and 2000 features. To assess the model fit and selection, we varied the noise variance σ (rows) and performed the SOFA factorization over a range of decomposition ranks with and without the use of guiding variables. **a**: Average root mean square error (RMSE) of true and predicted data averaged over 10 model initializations. **b-c**: Scatter plot of true guiding variable (Y) vs predicted guiding variable (Y) (**b**: binary; **c**: continuous) of one of the rank-5 models from **a**.

**Figure S2:**
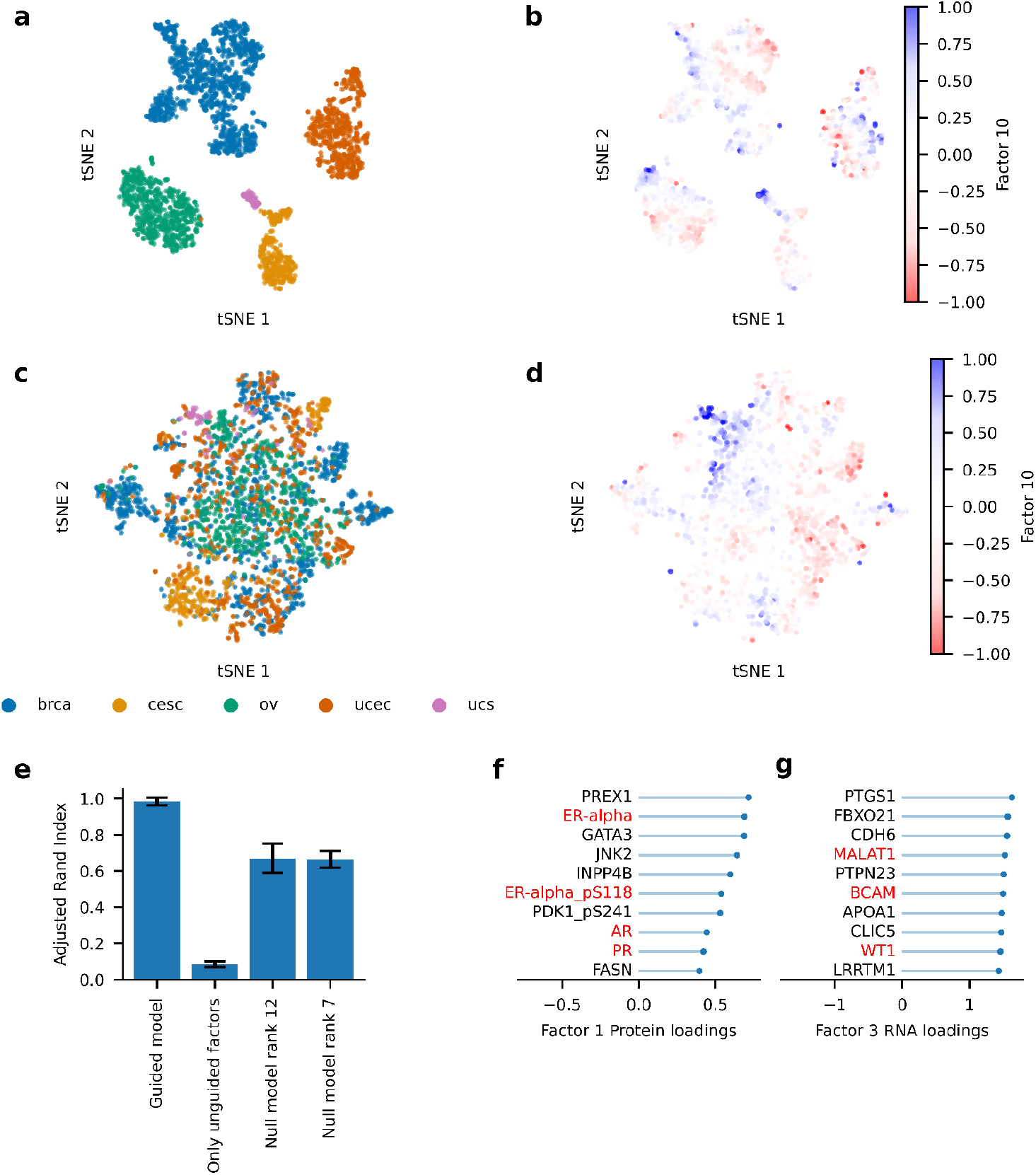
**a**: tSNE of all factors colored by cancer type. **b:** tSNE of all factors colored by Factor 10. **c**: tSNE of unguided factors 7-12 colored by cancer type. **d**: tSNE of unguided factors 7-12 colored by Factor 10. **e**: Adjusted Rand Index (ARI) for clusterings of all factors of the guided model, only unguided Factors 7-12, unguided null model of rank 12 and unguided null model of rank 7. The clusterings were compared to the true cancer type labels. A higher ARI suggests better overlap of the clustering with the cancer type labels. The ARI was computed for 10 independent model initializations. **f**: Highest protein loadings of Factor 1. **g**: Highest RNA loadings of Factor 3. brca: breast invasive carcinoma; cesc: Cervical squamous cell carcinoma and endocervical adenocarcinoma; ov: Ovarian serous cystadenocarcinoma; ucec: Uterine Corpus Endometrial Carcinoma; ucs: Uterine Carcinosarcoma

**Figure S3:**
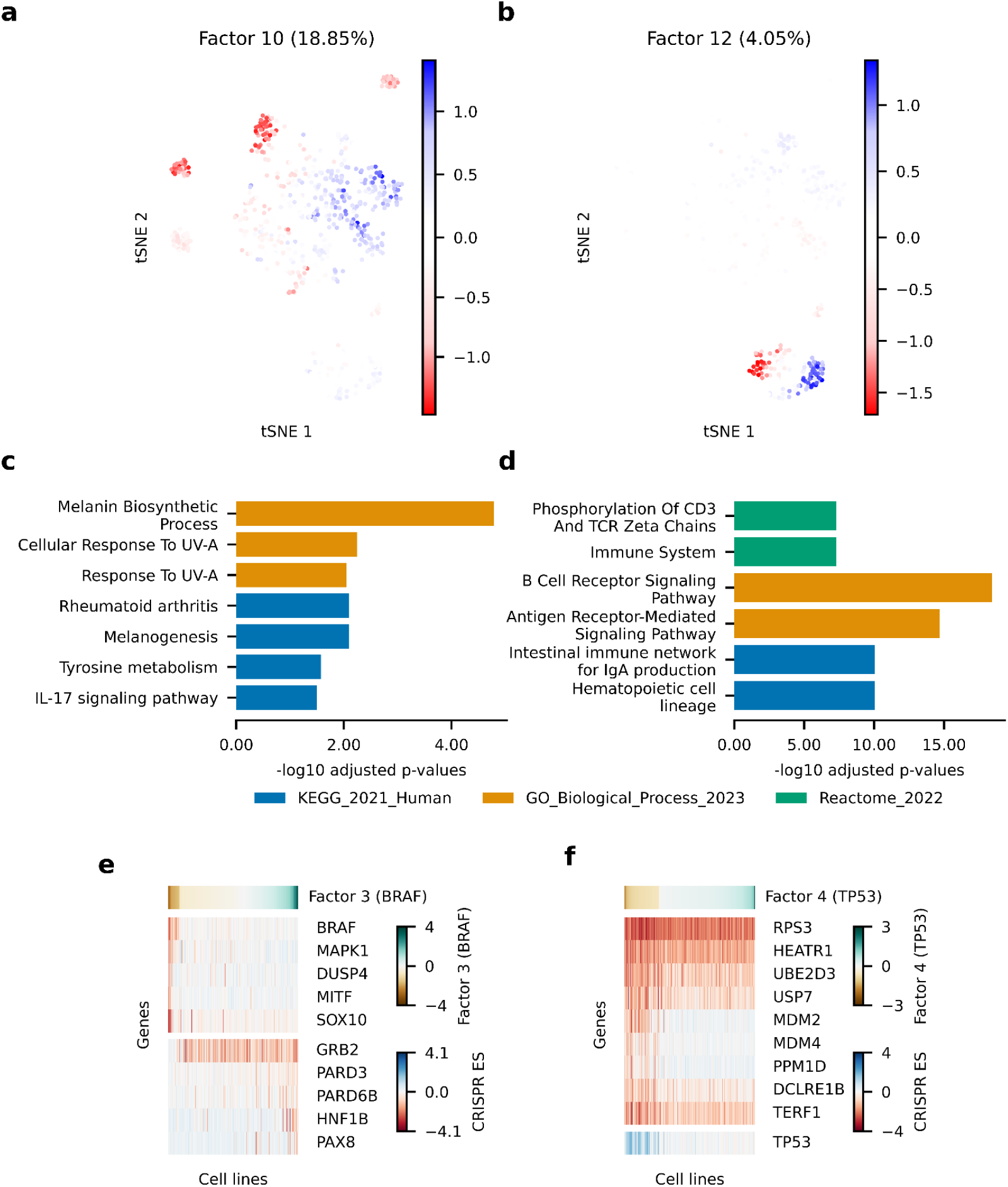
**a-b**: tSNE of all factors coloured by the values of factors 10 and 12. The fraction of explained variance is shown in parenthesis. **c**: Gene set overrepresentation analysis of highest RNA loadings of factor 3 (BRAF mutation). **d**: Gene set overrepresentation analysis of highest RNA loadings of factor 6 (hematopoietic lineage). **e**: Heatmap showing CRISPR essentiality scores (ES) of top 10 genes significantly associated with Factor 3 (BRAF mutation), tested using a linear model. The cell lines are ordered based on their values of Factor 3. **f**: Heatmap showing CRISPR essentiality scores (ES) of top 10 genes significantly associated with Factor 4 (TP53 mutation), tested using a linear model. The cell lines are ordered based on their values of Factor 4.

**Figure S4:**
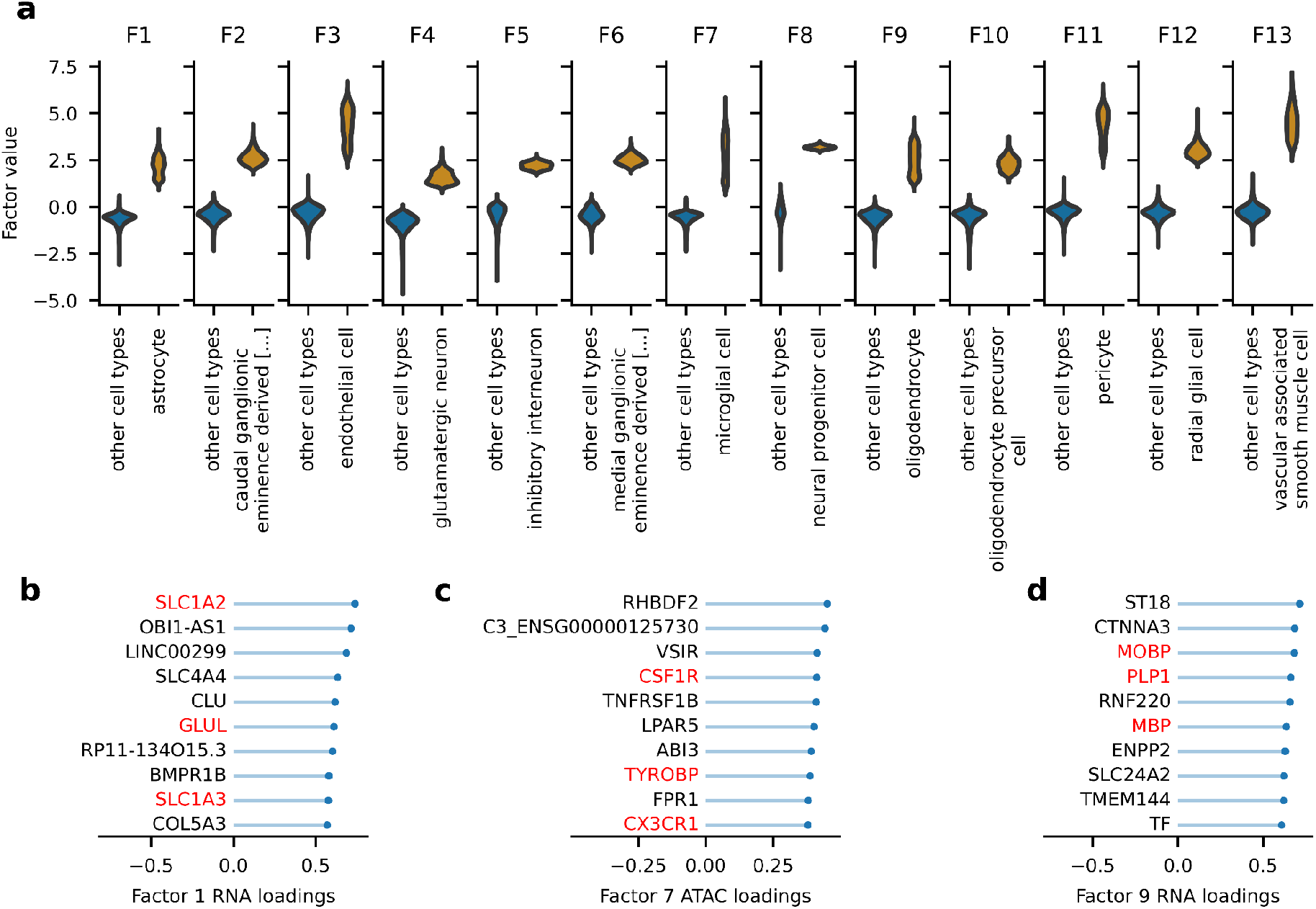
**a**: Violin plots of each guided factor vs their guiding cell type label. **b**: Highest transcriptomics (RNA) loadings of factor 1 (astrocytes). **c**: Highest ATAC loadings of factor 7 (microglial cells). **d**: Highest transcriptomics (RNA) loadings of factor 9 (oligodendrocytes).

**Figure S5:**
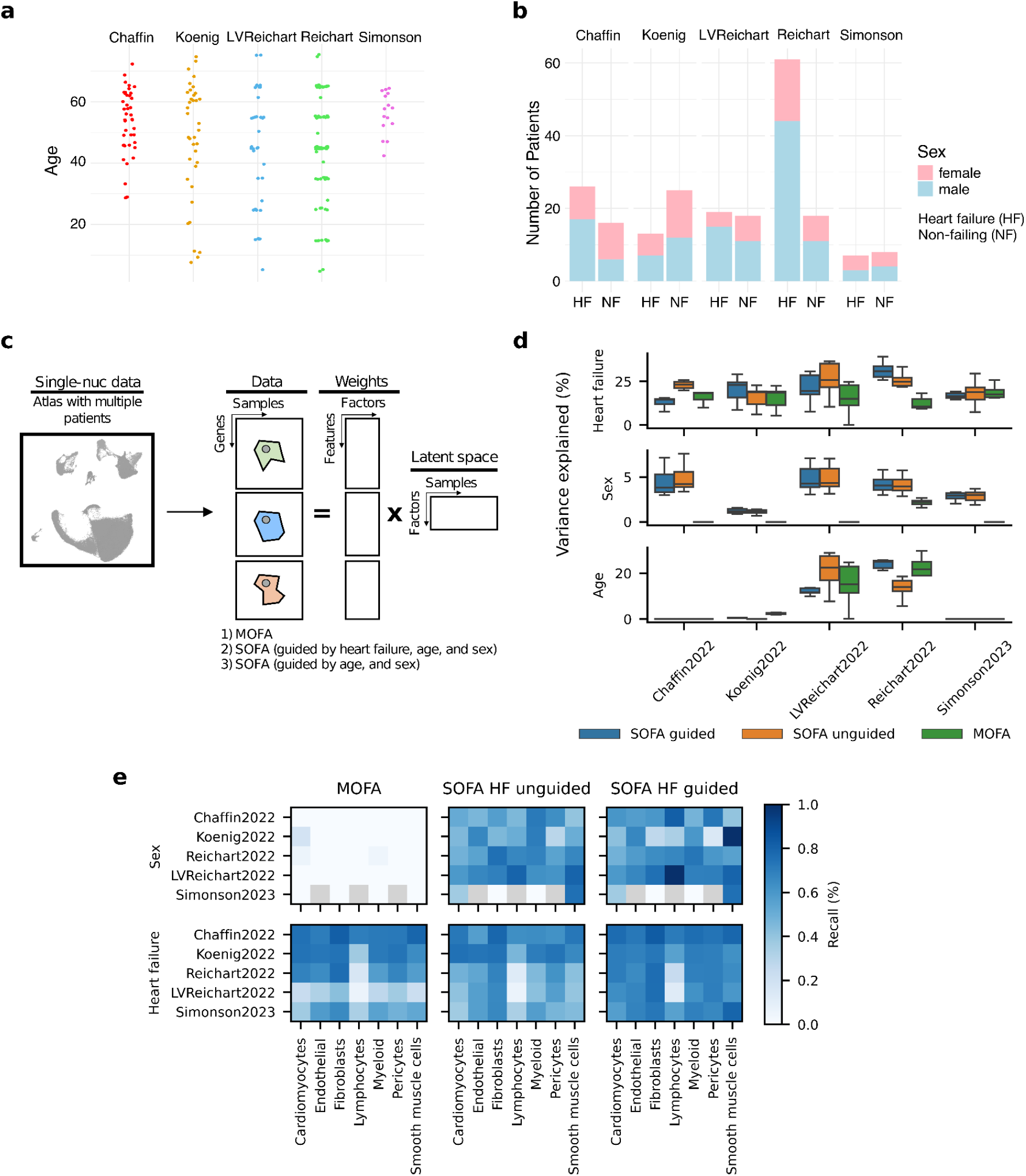
Comparison between SOFA and MOFA in the inference of multicellular programs from single-nuclei RNA-seq data: **a**: Age distribution across five patient cohorts. **b:** Sex distribution across five patient cohorts grouped by heart failure condition. **c:** Schematic of application of SOFA and MOFA for multicellular integration. Single-nuclei data is transformed into a multiview representation, where each view is the pseudobulk gene expression matrix of a cell-type. The latent space represents coordinated gene expression programs across cell-types or multicellular programs. **d:** Fraction of variance explained associated with heart failure, sex, and age across distinct datasets and models. **e:** Recall of differentially expressed genes for sex and heart failure conditions, based on gene sampling of equivalent size from the loading scores of the factors most strongly associated with each condition.

