## Supplementary Methods for "Semi-supervised Omics Factor Analysis (SOFA) disentangles known and latent sources of variation in multi-omic data"

#### 1 Setup

SOFA is based on group factor analysis (GFA), a multivariate statistical technique that describes linear relationships between groups of variables that were measured on a common set of objects or *samples*. Each group represents a set of particularly related variables (Klami et al., 2014). Multi-Omics Factor Analysis (MOFA, Argelaguet et al. (2018)) is an adaptation of this idea to multi-omics data in biology, where a series of biological specimen are assayed using multiple 'omics technologies, also called *views*. Each view corresponds to a group of variables. A typical objective in applications of these methods is to find a lower dimensional representation of the data that captures correlation structures that are shared across the views, as well as correlation structures among variables within, and unique to, individual views. Mathematically, these correlation structures are represented by factors (see below).

SOFA extends GFA by enabling the analyst to "guide" a subset of the factors by explicitly known guide variables for the samples, while others are assumed to be unobserved, or *latent*. The parameters of the model are inferred using black box variational inference (Ranganath et al., 2014).

We consider data $\{\mathbf{X}^1, \ldots, \mathbf{X}^M, \mathbf{Y}^1, \ldots, \mathbf{Y}^C\}$, where $\mathbf{X}^m \in \mathbb{R}^{N \times D^m}$ is the matrix of measurements for the $m$-th view, for $N$ samples with $D^m$ features each, and $\mathbf{Y}^c \in \mathbb{R}^N$ is the vector of measurements of the $c$-th known guide variable. The samples are the same, and aligned, across modalities and variables. There is no assumption of relationships or alignments between the features from the different views. SOFA finds a reduced-dimensional representation of these data using $K \leq \min\{D^1, \ldots, D^M\}$ factors:

$$\mathbf{X}^m = \mathbf{Z} \times \mathbf{W}^m + \boldsymbol{\varepsilon}^m, \qquad m = 1, \ldots, M \tag{1}$$

with the latent factors $\mathbf{Z} \in \mathbb{R}^{N \times K}$, the loadings $\mathbf{W}^{\mathbf{m}} \in \mathbb{R}^{K \times D^m}$, and the residuals $\boldsymbol{\varepsilon}^m \in \mathbb{R}^{N \times D^m}$. We model the residuals as independent random variables from Normal distributions with feature- and view-specific standard deviation $\sigma_j^m$.

$$\varepsilon_{ij}^m \sim \mathcal{N}(0, \sigma_j^m) \qquad \forall \qquad i = 1, \ldots, N \qquad j = 1, \ldots, D^m. \tag{2}$$

This implies for the data

$$x_{ij}^m \sim \mathcal{N}\left(\sum_{k=1}^{K} z_{ik} w_{kj}^m, \sigma_j^m\right). \tag{3}$$

We set no priors on the standard deviation parameters $\sigma_j^m$.

For model inference, we first set aside the variables $\mathbf{Y}^c$ (we bring them back in Section 2). We posit standard Normal priors on the factors,

$$z_{ik} \sim \mathcal{N}(0,1), \tag{4}$$

and hierarchical horseshoe priors (Carvalho et al., 2009) on the loadings $\mathbf{W^m}$. This assumption encourages sparsity, facilitates model interpretability, and is justified by the observation that many useful models of biological systems use sparse sets of relationships between the system components. To this end, we introduce the parameters $\lambda_{kj}^m$, $\delta_k^m$ and $\tau^m$ and assume half Cauchy priors for them:

$$\begin{aligned} \tau^m &\sim \mathcal{C}^+(0,1) \\ \delta_k^m &\sim \mathcal{C}^+(0,1) \\ \lambda_{kj}^m &\sim \mathcal{C}^+(0,1). \end{aligned} \tag{5}$$

The parameter $\tau^m$ is used to generate view-level sparsity (i.e., whole views can be deactivated), $\delta_k^m$ generates factor-level sparsity (i.e., whole factors can be deactivated within individual views), and $\lambda_{kj}^m$ generates feature-level sparsity (i.e., individual features can be deactivated within individual factors and views). Then,

$$w_{kj}^m \sim \mathcal{N}(0, \lambda_{kj}^m \delta_k^m \tau^m). \tag{6}$$

#### 2 Adding factor guidance

SOFA associates the first $C$ factors with the guide variables $\mathbf{Y}^1, \ldots, \mathbf{Y}^C$ by linear regression. Intuitively, this constrains this set of "guided" factors to explain sources of variation in $\mathbf{X}^1, \ldots, \mathbf{X}^M$ with factors that are highly correlated with the variables $\mathbf{Y}^1, \ldots, \mathbf{Y}^C$. Conversely, the remaining unguided factors are free to capture further sources of variation in $\mathbf{X}^1, \ldots, \mathbf{X}^M$ that are less correlated with the guide variables.

Thus, we posit a generalized linear model (Nelder and Wedderburn, 1972)

$$f\left(\mathbb{E}[y_i^c]\right) = \alpha_c + \beta_c z_{ic}, \tag{7}$$

where $f$ is a link function, and we assume a standard Normal prior on the offset and slope coefficients,

$$\alpha_c, \beta_c \sim \mathcal{N}(0,1). \tag{8}$$

SOFA supports continuous and categorical response variables by supporting Normal, Bernoulli and Categorical likelihood functions. For real-valued continuous variables, link function and likelihood are

$$f(y) = y \tag{9}$$

$$y_i^c \sim \mathcal{N}\left(\alpha_c + \beta_c z_{ic}, \kappa_c^2\right) \tag{10}$$

where $\kappa_c^2$ denotes the noise variance. For binary variables,

$$f(p) = \log \frac{p}{1-p} \tag{11}$$
$$y_i^c \sim \text{Bernoulli}\left(f^{-1}(\alpha_c + \beta_c z_{ic})\right). \tag{12}$$

For a categorical variable with $L$ levels,

$$f^{-1}(x_1, \ldots, x_L) = \frac{1}{\sum_{l'=1}^{L} e^{x_{l'}}} (e^{x_1}, \ldots, e^{x_L}) \tag{13}$$
$$y_i^c \sim \text{Categorical}\left(f^{-1}(\boldsymbol{\alpha}_c + \boldsymbol{\beta}_c z_{ic})\right), \tag{14}$$

where $\boldsymbol{\alpha}_c, \boldsymbol{\beta}_c$ are vectors of length $L$ whose elements come from the prior (8).

### 3 Inference

For the implementation of SOFA, we used the probabilistic programming framework Pyro (Bingham et al., 2019). In the following, we describe the inference for the case that all $\mathbf{Y}_c$ are normal, as in Eqn. (10). The likelihood is then

$$\begin{aligned}
p(\mathbf{X}, \mathbf{Y}, \mathbf{Z}, \mathbf{W}, \boldsymbol{\lambda}, \boldsymbol{\beta}, \boldsymbol{\tau}, \boldsymbol{\delta}) = \\
&\prod_{m=1}^{M} \prod_{i=1}^{N} \prod_{j=1}^{D^m} \mathcal{N}(x_{ij}^m \mid \sum_{k=1}^{K} z_{ik} w_{kj}^m \lambda_{kj}^m \delta_k^m \tau^m, \sigma_j^m) \\
&\prod_{i=1}^{N} \prod_{c=1}^{C} \mathcal{N}(y_i^c \mid \alpha_c + z_{ic}\beta_c, \kappa_c^2) \\
&\prod_{i=1}^{N} \prod_{k=1}^{K} \mathcal{N}(z_{ik} \mid 0, 1) \\
&\prod_{m=1}^{M} \prod_{j=1}^{D^m} \prod_{k=1}^{K} \mathcal{N}(w_{kj}^m \mid 0, 1)\, \mathcal{C}^+(\lambda_{kj}^m \mid 0, 1)\, \mathcal{C}^+(\delta_k^m \mid 0, 1)\, \mathcal{C}^+(\tau^m \mid 0, 1) \\
&\prod_{c=1}^{C} \mathcal{N}(\beta_c \mid 0, 1)
\end{aligned} \tag{15}$$

We make use of the mean field assumption (Blei et al., 2017) and introduce a factorized family of parameterized distributions to approximate the intractable posterior,

$$q_\phi(\Theta) = \prod_{\theta \in \Theta} q_\phi(\theta), \tag{16}$$

where $\Theta$ denotes the collection of inferred parameters $\{\mathbf{Z}, \mathbf{W}, \boldsymbol{\lambda}, \boldsymbol{\beta}, \boldsymbol{\tau}, \boldsymbol{\delta}\}$ and $\phi$ the parameters of the variational distribution $q(\cdot)$. For the variational distribution, we posit the following form using Normal and log-Normal distributions:

$$
\begin{aligned}
q(\mathbf{Z}, \mathbf{W}, \boldsymbol{\lambda}, \boldsymbol{\beta}, \boldsymbol{\tau}, \boldsymbol{\delta}) = &\prod_{i=1}^{N} \prod_{k=1}^{K} \mathcal{N}(z_{ik} \mid m_{ik}, s_{ik}) \\
&\prod_{m=1}^{M} \prod_{j=1}^{D^m} \prod_{k=1}^{K} \mathcal{N}(w_{kj}^m \mid m_{kj}^m, s_{kj}^m) \\
&\prod_{m=1}^{M} \prod_{j=1}^{D^m} \prod_{k=1}^{K} \log\mathcal{N}(\lambda_{kj}^m \mid m_{kj}^m, s_{kj}^m) \\
&\prod_{m=1}^{M} \prod_{k=1}^{K} \log\mathcal{N}(\delta_k^m \mid m_k^m, s_k^m) \\
&\prod_{m=1}^{M} \log\mathcal{N}(\tau^m \mid m^m, s^m) \\
&\prod_{c=1}^{C} \mathcal{N}(\beta_c \mid m_c, s_c)
\end{aligned}
\tag{17}
$$

We optimize the evidence lower bound (ELBO) with respect to the variational parameters $\phi = \{\mathbf{m}, \mathbf{s}\}$. $\mathbf{m}$ and $\mathbf{s}$ are the means and standard deviations of the factors of the variational distribution in Eqn. (17), where we dropped the indices for notational simplicity. The ELBO is given by:

$$
\text{ELBO}(\phi) = \mathbb{E}_{q_\phi}[\log p(\mathbf{X}, \mathbf{Y}, \boldsymbol{\Theta}) - \log q_\phi(\boldsymbol{\Theta})]. \tag{18}
$$

The ELBO is optimized using stochastic variational inference by taking Monte Carlo samples from the variational distribution and computing unbiased but noisy gradients with respect to the variational parameters from the ELBO (Hoffman et al., 2013; Ranganath et al., 2014). The noise scales $\sigma$ for $\mathbf{X}$ and $\kappa^2$ for $\mathbf{Y}$ are fit via maximum likelihood, and no prior distributions are imposed on them. Optimization is stopped when the change in ELBO is smaller than a user defined threshold over the last 2000 iterations.

If necessary for large datasets, e.g., in applications to single-cell RNA-sequencing data, we compute the gradients over subsamples of batches of the observations.
